# Did *Caenorhabditis* nematodes recycle transposons to fight pathogens?

**DOI:** 10.1101/2022.01.28.478163

**Authors:** Zixin Li, Christian Rödelsperger

## Abstract

Animal genomes consist largely of sequences derived from transposons which were previously considered as junk DNA and active transposons can even be deleterious for organismal fitness. Nevertheless, the huge pool of transposon-derived sequences also forms the raw material for molecular innovations. Here, we follow up on the incidental finding of a transposon-derived DNA binding domain in a subset of F-box genes in *Caenorhabditis elegans*. Based on phylogenetic analysis, we show that a single gene fusion followed by individual losses explains most members of this novel gene family. Phylogenomic data of available *Caenorhabditis* genomes allowed us to trace this fusion event to the ancestor of the Elegans group. Additional homology searches suggest endogenous *Mariner* transposons as the likely source of the coopted sequence. Further bioinformatic characterization of different F-box families by Gene Ontology analysis, gene module comparisons, and literature research identified first evidence that some F-box genes might be involved in innate immunity, as it had been proposed previously based on adaptive signatures of molecular evolution. Specifically, the F-box gene *fbxa-158* contains one of the transposon-derived domains and was shown to interact with the components of the intracellular pathogen response machinery targeting Microsporidia and viruses. Thus, cooption of transposon-derived sequences likely contributed to the adaptive evolution of the F-box superfamily in *Caenorhabditis* nematodes.

**Significance statement:** When considering transposons as genomic junk or selfish elements that are detrimental for organismal fitness, we often neglect the potential of transposon-derived sequences as a source of molecular innovation. Here, we characterize a case where a transposon-derived sequence has been coopted by one of the largest and fastest evolving gene superfamilies in *Caenorhabditis* nematodes which include the model organism *C. elegans*. The resulting chimeric gene family has been stably maintained for about 20 million years and bioinformatic analysis reveals first evidence for potential functions in innate immunity. Thus, strong evolutionary pressure might have forced *Caenorhabditis* nematodes to recycle transposons in order to fight pathogens.

## Introduction

The genome of the nematode *Caenorhabditis elegans* was the first to be sequenced from a multicellular organism (C. elegans Sequencing Consortium 1998). Subsequent genome projects of parasitic and freeliving nematodes revealed that they typically have quite compact genomes. One potential explanation for this might be that they are more effective in controlling the expansion of transposons (Rödelsperger et al. 2013). Thus, only 12% of the *C. elegans* genome consists of repeats (C. elegans Sequencing Consortium 1998). Despite being one of the most extensively studied model organisms, more than 40% of *C. elegans* genes are still without any functional annotation and more than 60% of genes have no phenotypic association (Petersen et al. 2015). This lack of biological understanding might be especially prevalent in large gene families where genes may act redundantly. Fifteen years ago, James H. Thomas published a study of two large gene families, the BTB family and the F-box superfamily (Thomas 2006). Both families are thought to represent substrate-specific adapter proteins in ubiquitin ligase complexes that are responsible for protein degradation (Moon et al. 2004). Analysis of nonsynonymous and synonymous substitutions between paralogs revealed widespread evidence of positive selection. Together with the finding of accelerated birth and death rates and the overall high diversity of these two large gene families, Thomas proposed that they might be involved in innate immunity. The observed signature of molecular evolution would hereby reflect a host-pathogen arms race and is similar to patterns in other large gene families like C-type lectins which are known to be involved in innate immunity (Kogelberg & Feizi 2001; Pees et al. 2016). Based on the combinations of protein domains, Thomas defined three F-box families, A1, A2, and B (Fig. 1). The latter was defined by a combination of an F-box and an FBA2 domain. Family A1 members have an F-box and an FTH domain. Family A2 was found to possess an additional domain that is related to the DNA binding domain of a *Mariner* transposase (Thomas 2006). However, neither him nor anyone else ever followed up on this incidental observation. Even though repetitive sequences were considered as junk and transposon activity can be deleterious, in rare cases, transposable elements can be a source of molecular innovation, as they were shown to rewire transcriptional networks and their sequences can be coopted to form novel gene families (Toll-Riera et al. 2009; Kunarso et al. 2010; Cosby et al. 2021; Chuong et al. 2016; Carelli et al. 2022). Therefore, we decided to revisit Thomas’ finding, to characterize the potential origin of this new gene family, and to search for experimental evidence that would support his innate immunity hypothesis.

**Fig. 1.**
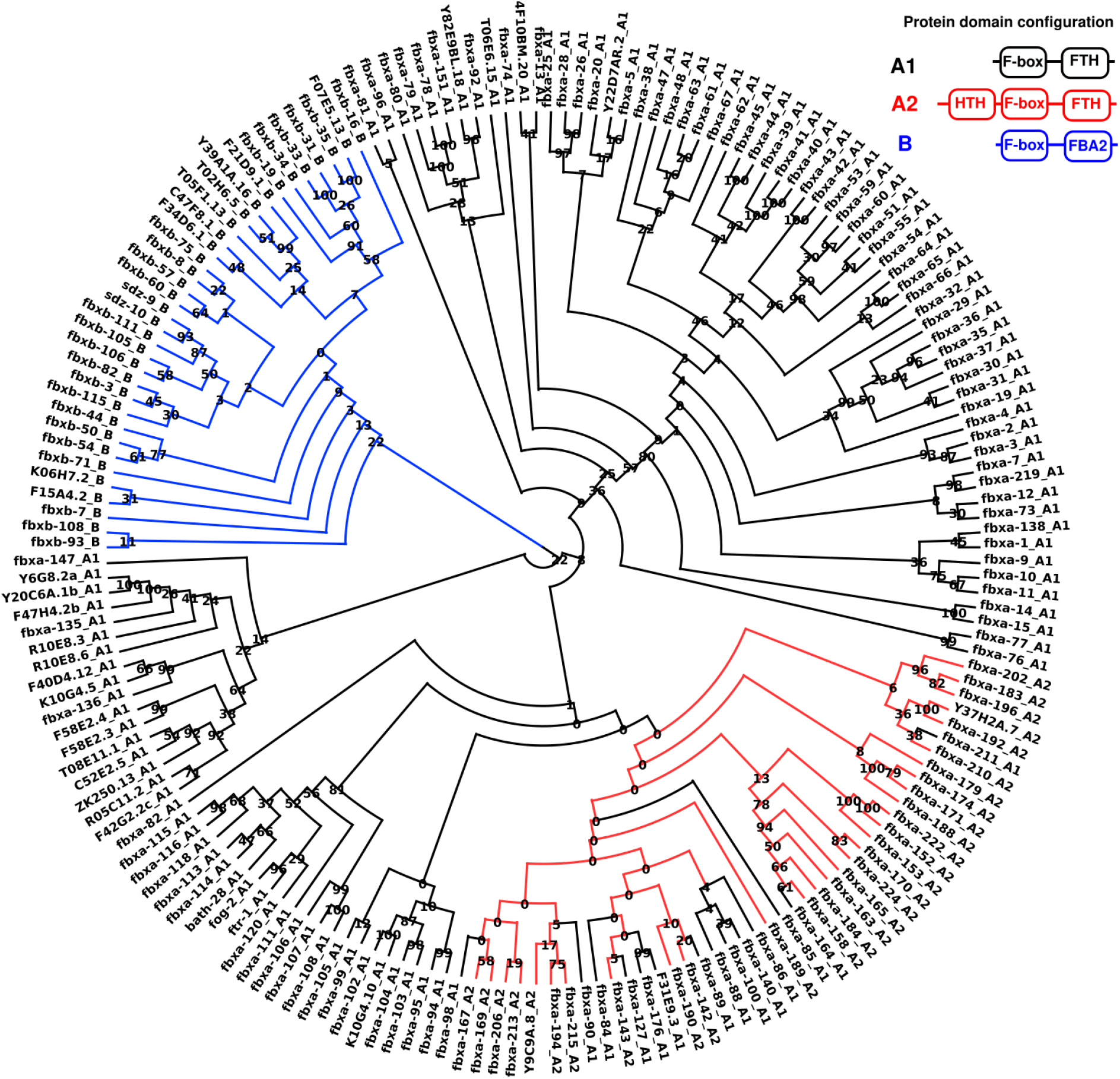
A single gene fusion with subsequent losses explains most A2 genes. An alignment of all F-box proteins from *C. elegans* was used to compute a maximum-likelihood tree. The legend shows the color code for all three gene families and a schematic of their protein domain configurations. Numbers at internal branches indicate bootstrap support values (100 pseudoreplicates). Most A2 family members fall into a subtree within the larger A1 family. Individual genes within the A2 subtree seem to have lost the HTH domain.

## Results

### A single gene fusion explains the majority of A2 family members

To characterize F-box genes in *C. elegans*, we identified gene family members based on protein domain analysis using the definitions from Thomas (2006). Thereby, the A2 family was defined by the presence of an additional Helix-Turn-Helix (HTH) domain (PF17906) that forms the DNA binding domain of the *Mos1* transposase in *Drosophila* (Richardson et al. 2009). In total, we found 118 members of family A1, 30 members of family A2, and 34 members of family B (Supplemental Table S1). To test whether the A2 family has a single or multiple origins, we performed a phylogenetic analysis of all *C. elegans* members of these three gene families (Fig. 1, Supplementary Fig. S1). While family B is found in a completely monophyletic clade, the pattern of A1 is more patchy. Under the assumption that A2 members date back to a single evolutionary event, multiple independent losses have to be postulated. Alternatively, HTH domains could have been translocated across members of different subtrees. However, even if the phylogenetic analysis cannot conclusively show a single evolutionary origin of the A2 family, it still seems plausible that at least the majority of the A2 genes are derived from such an event. This is further supported by individual loss of other domains. For example, *fbxa-197* has lost the F-box domain and Y37H2A.12 has no FTH domain (Supplemental Table S1). Thus, the evolution of most A2 family members can be explained by an ancestral gene fusion followed by individual domain loss events.

### The A2 family is only found in *Caenorhabditis* species of the Elegans group

While the phylogenetic analysis supports that a single fusion between an HTH domain and an F-box gene of the A1 family gave rise to a new gene family, it is not clear when this fusion occurred. To date the origin within the evolutionary history of nematodes, we screened phylogenomic data of other *Caenorhabditis* species and multiple outgroups for the presence of members of the A1, A2, and B families (Fig. 2). This revealed that all three F-box families represent *Caenorhabditis*-specific molecular innovations. Furthermore, the three families only explain a fraction of all F-box genes. These observations suggest that a global reclassification of the F-box superfamily using a broad taxonomic sampling might be needed. The A2 family is only present in the Elegans group and likely originated in its ancestor. Keeping in mind the difficulties to estimate divergence times in nematodes, this would translate into an approximate age of roughly 20 million years for the A2 family (Cutter 2008). This shows that the A2 family survived multiple speciation events and is therefore evolutionary stable.

**Fig. 2.**
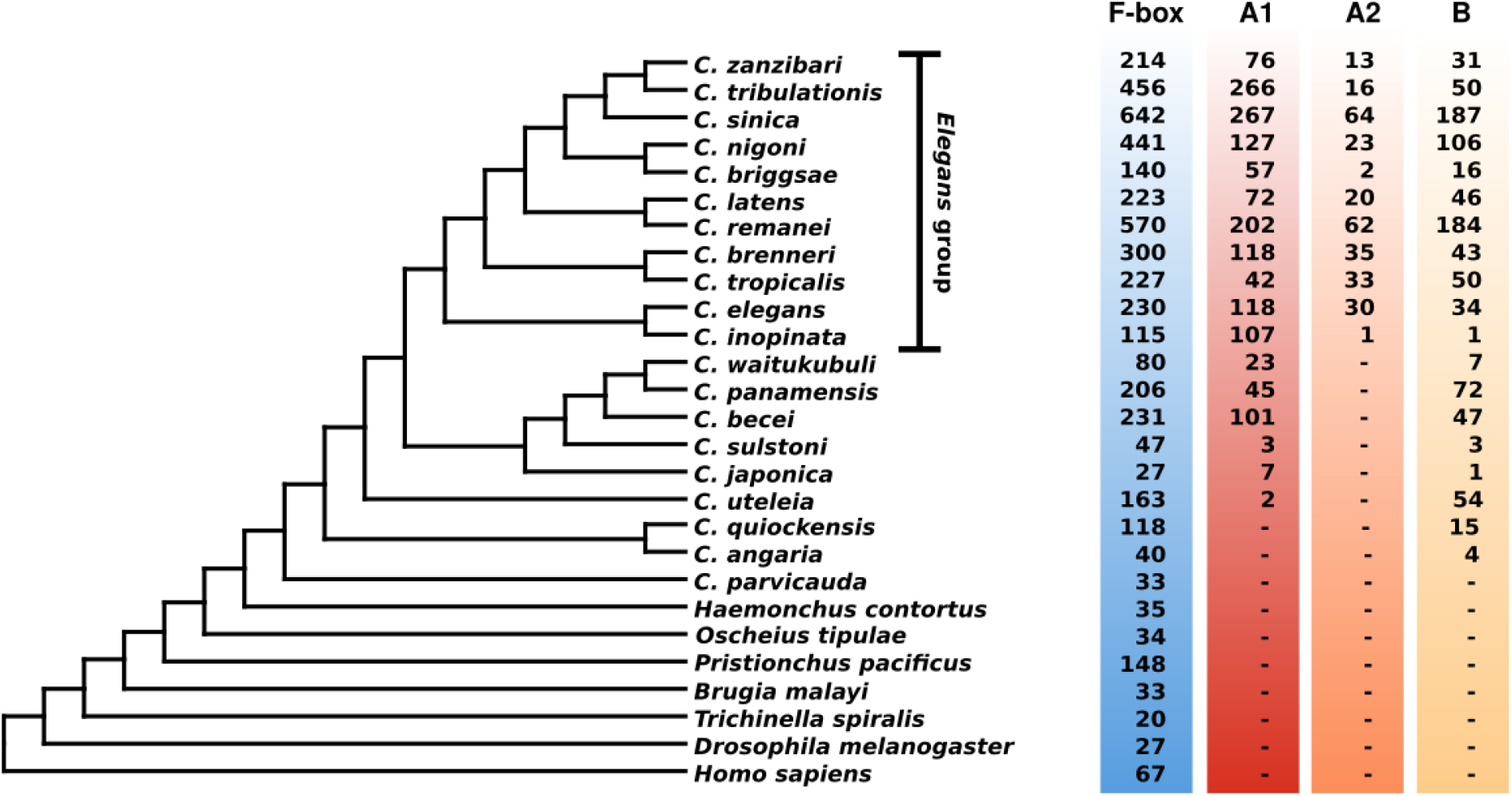
The A2 family originated in the ancestor of the Elegans group. Comparative genomic data from 27 species was used to date the origin of the A2 family. The schematic phylogeny was redrawn based on previous phylogenomic analyses (Stevens et al. 2019; Smythe et al. 2019). For each species, the total number of F-box containing genes is shown, as well as the members of all three families as defined by Thomas (2006). Only *Caenorhabditis* species of the Elegans group have A2 members, which indicates that the gene fusion occurred in the ancestor of the Elegans group.

### Phylogenetic analysis points towards an origin of the HTH domain from an endogenous transposon

The homology with the HTH domain (PF17906) from the Pfam database is already a strong indication that the fused sequence is of transposon origin. However, it is not clear whether the A2 ancestor received this sequence directly from an endogenous or horizontally transferred transposon or from an intermediate gene family that was the original recipient of the HTH domain. Further analysis of *Caenorhabditis* genes with HTH domains showed that this domain typically occurs either in combination with a transposase (PF01359) or within an F-box context (either with an FTH, an F-box domain, or with both). These observations rule out the possibility of a secondary transition. Next, we searched for the most closely related sequences in species outside the Elegans group and in non-coding regions of the *C. elegans* genome. Phylogenetic analysis showed that one of the most closely related sequences corresponds to an intronic region that was annotated as a Mariner element in the *C. elegans* genome (Fig. 3, Supplementary Figure S2,S3). This suggests that the HTH domain is likely derived from an endogenous transposon. However, the overall tree topology is only poorly supported and we cannot completely rule out the possibility of an origin from a horizontally transferred transposon (Palazzo et al. 2021).

**Fig. 3.**
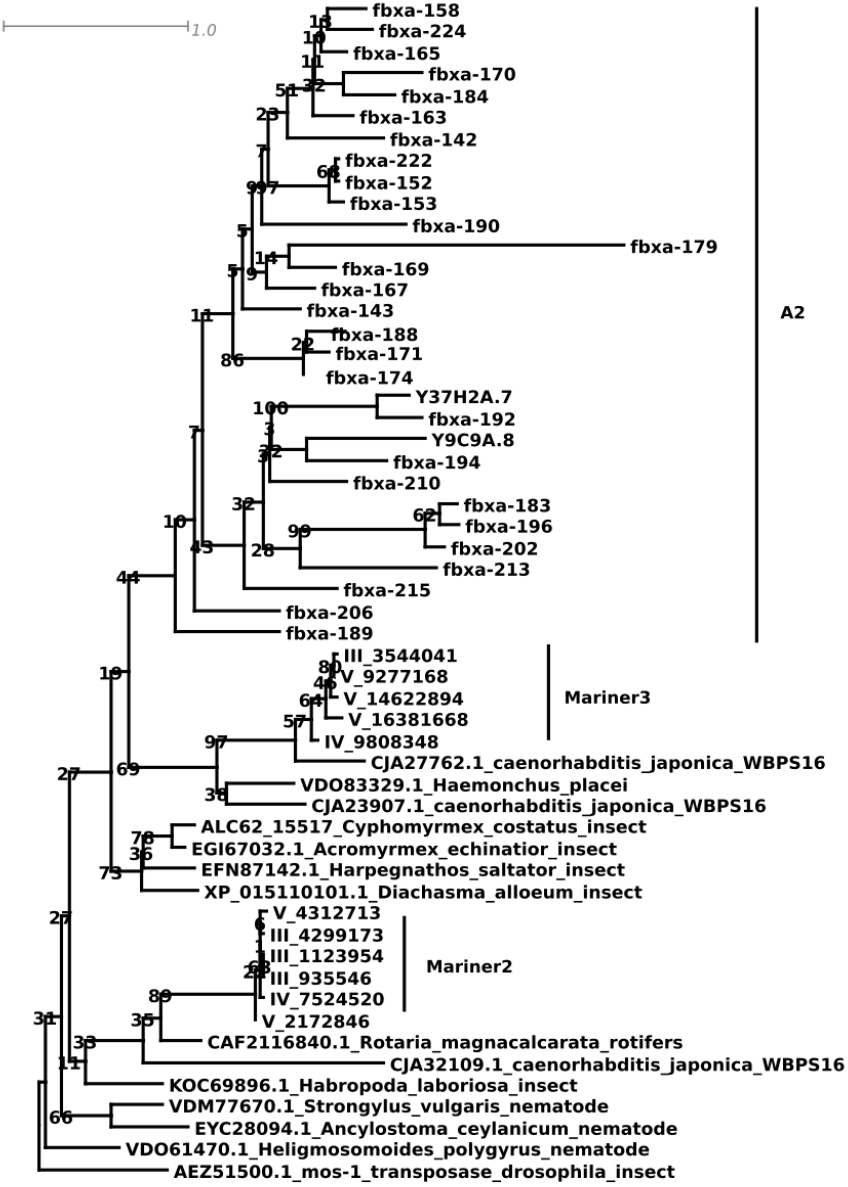
The likely donor of the HTH domain was an endogenous transposon. Phylogenetic analysis was performed for the HTH domains from F-box genes of the A2 family, their most closely-related sequences outside the Elegans group, and non-coding hits in the *C. elegans* genome. One of the most closely related sequences corresponds to an intronic *Mariner* element in *C. elegans* (Supplementary Fig. S2, S3) which suggests that the HTH domain was derived from an endogenous transposon.

### Analysis of functional and expression data indicates towards roles in immune response

To characterize biological functions of the F-box gene families, we performed a gene ontology analysis using the David platform (Huang et al. 2009). Only the largest family A1 showed a single significant enrichment in the biological process of innate immune response (GO:0045087, Benjamini corrected *P* = 1.2 x 10^-5^). No other significantly enriched terms were detected for either family A1 or any of the other two gene families. Next, we tested the different gene families for overlap with 209 coexpression modules that were defined by large-scale meta-analysis of more than thousand expression profiling experiments (Cary et al. 2020). Family A1 was most significantly enriched in modules 55 and 200 and family A2 was significantly enriched in module 32 (Benjamini-Hochberg corrected *P* < 0.001). Comparison between the weight vectors of the gene modules and expression fold changes from individual studies showed that these modules reflect expression changes in worms upon exposure to either bacterial or viral pathogens (Estes et al. 2010; Sarkies et al. 2013). Family B was enriched in module 116 (Benjamini-Hochberg corrected *P* < 0.001) which reflects expression changes during embryonic development (Baugh et al. 2005). Thus, both analyses support that family A members could play a role in innate immunity.

### The A2 gene *fbxa-158* interacts with components of the intracellular pathogen response

Finally, we screened biomedical literature for detailed experimental studies of individual F-box genes. In family B, only *fbxb-3* was found in an RNAi screen for altered embryonic development (Du et al. 2015). Together with the results of the gene module analysis, this potentially suggests a trend of family B members towards being involved in embryonic development. Out of the A1 family, the genes *fog-2* and its paralog *ftr-1* have been characterized previously. Mutations in *fog-2* transform hermaphrodites into females that are not capable of self-fertilization (Schedl & Kimble 1988). *fog-2* arose recently via gene duplication in *C. elegans* and has no one-to-one ortholog in *C. briggsae* (Nayak et al. 2005). The only member of family A2 which has been experimentally studied is *fbxa-158. fbxa-158* was found to be differentially expressed in response to intracellular pathogens such as microsporidia and viruses (Bakowski et al. 2014; Sarkies et al. 2013). This intracellular pathogen response pathway was later shown to promote proteostasis during infection (Reddy et al. 2017). Finally, co-immunopreciptation, RNAi and gene editing experiments showed that *fbxa-158* interacts with components of the intracellular pathogen response pathway (Panek et al. 2020). Thus, *fbxa-158* is a member of the F-box A2 family with experimental evidence for a potential role in innate immunity.

## Discussion

New genes are introduced into genomes by diverse mechanisms including duplication, *de novo* formation, and horizontal gene transfer (Rödelsperger et al. 2019). Here, we describe a particular example, where the DNA binding domain of a transposon was coopted to form a new chimeric family of F-box genes. This highlights the role of repetitive and transposable elements as a reservoir for molecular innovations (Toll-Riera et al. 2009; Cosby et al. 2021; Bourque et al. 2018; Kapusta et al. 2013). The F-box superfamily might be specifically predisposed for such events as it seems to underlie some force of divergent selection. The evidence of positive selection and gene turnover, as observed by James H. Thomas, has been further supported by the analysis of genomic variation across natural isolates (Maydan et al. 2010; Volkers et al. 2013; Ma et al. 2021). What is driving this rapid evolution is still not clear. Fifteen years after the proposition of the innate immunity hypothesis, available experimental data is still insufficient to either confirm or refute the idea that the molecular evolution of F-box proteins reflects an arms race between host and pathogens. It is very plausible that F-box proteins may play key roles during the response to pathogens as the importance of protein degradation pathways during immune response is well recognized (Reddy et al. 2017; Gallotta et al. 2020; Garcia-Sanchez et al. 2021). In addition, the finding of *fbxa-158* to be involved in the immune response is the first experimental support for the innate immunity hypothesis, but it is by far not enough to prove it. Knockouts of *fbxa-158* also affect thermotolerance, which is another phenotype that is tightly linked to proteostasis (Panek et al. 2020). Thus, F-box proteins in general also including the A2 family could be involved in protein degradation at a variety of biological processes. Another open question is what is the role of a DNA binding domain. We are tempted to speculate that it could possibly recognize viral DNA. Another possible explanation could be that it contributes to the control of transposon activity. Consistent with this, multiple F-box genes including the A2 member *fbx-196* have been recently found to be expressed during spermatogenesis (Rödelsperger et al. 2021).

One major problem in the investigation of large gene families might be genetic redundancy. However, with the advent of efficient genome editing techniques systematic functional characterization of partial or complete gene families may be feasible (Han et al. 2022; Baker et al. 2021; Sieriebriennikov et al. 2020). Yet, to get a broader understanding of gene functions, we also need a concerted effort to study genes in a less biased manner. Our analysis of biomedical literature found 82 studies on *fog-2*, the best characterized F-box gene in *C. elegans.* The second best studied is its paralog *ftr-1* with two papers while almost all other F-box genes were never mentioned in titles or abstracts. The F-box superfamily is by far not the only understudied gene class in *C. elegans*. We recently reported an association of two uncharacterized gene families with spermatogenesis related tissues (Rödelsperger et al. 2021). Another factor that limits the biological understanding of individual gene function is the environmental influence. For decades, *C. elegans* was studied under standardized laboratory conditions on an artificial food source. Only recently, the field has started to investigate its biology under more natural and diverse contexts (Dirksen et al. 2016, 2020). Thus, there is hope that in fifteen years from now, we can answer the question whether *Caenorhabditis* nematodes recycled transposons to fight pathogens.

## Methods

### Protein domain annotation and phylogenetic analysis

Comparative genomic data of several nematode species was downloaded from WormBase Parasite (version: WBPS16) (Howe et al. 2017). For *Pristionchus pacificus,* we used the latest gene annotation (version: El Paco gene annotation V3) (Athanasouli et al. 2020). Protein sequence for *Homo sapiens* and *Drosophila melanogaster* were downloaded from the Ensembl (release-93) and Ensembl Metazoa (release-40) ftp servers. In case of multiple isoforms per gene, the longest protein was chosen as representative isoform. Protein domain annotation was done using the hmmsearch program of the HMMer package (version 3.3, option -e 0.001) by searching against the Pfam database Pfam-A.hmm (version 3.1b2, Febrary 2015) (Mistry et al. 2013, 2021). F-box containing genes were defined on the presence of one of the domains PF00646 (F-box), PF12937 (F-box-like), PF13013 (F-box-like_2), PF15966 (F-box_4), PF18511 (F-box_5). The FTH and FBA2 domain were defined by the Pfam profiles PF01827 and PF07735, respectively. For phylogenetic analysis, F-box genes of all three families in *C. elegans* were aligned with the MUSCLE software (version 3.8.31) and the program raxml was run to compute a maximum-likelihood tree (version: 8.2.11, options: -m PROTGAMMAILG -f a) using 100 pseudoreplicates to compute bootstrap support values (Edgar 2004; Stamatakis 2014). To screen for homologous HTH domains, we performed a BLASTP search for *C. elegans* HTH domains against the NCBI nr database excluding any *Caenorhabditis* species and a BLASTP search against *Caenorhabditis* species outside of the Elegans group on WormBase ParaSite. Homologous non-coding sequences were identified by TBLASTN searches against the *C. elegans* genome (version: 2.10.1+, options: -max_target_seqs 2 -evalue 0.001). Phylogenetic analysis was performed as described above. Visualization of the multiple alignment was done using seaview (version 4.7 with default options) (Gouy et al. 2010).

### Functional characterization

Gene Ontology analysis was performed by the David Webtool using default options (Huang et al. 2009). Specifically, the Functional Annotation Chart functionally was run by testing all three F-box gene families against the Ontologies for Biological Process (GOTERM_BP_DIRECT), Cellular Component (GOTERM_CC_DIRECT), and Molecular Function (GOTERM_MF_DIRECT). In addition, we tested the gene sets from all three F-box gene families for significant overlaps with 209 coexpression modules that were obtained from an independent component analysis of more than thousand gene expression profiling studies (Cary et al. 2020). Comparison between the weight vector of the significant modules with existing expression data was done by the Tool 2 of the genemodules.org website. Precisely, we used the highest positive signed variance explained (SVE) measure to associate a module with a gene expression study (Cary et al. 2020). For literature research, the get_pubmed_ids function of the R package “easyPubMed” was used to automatically query the PubMed database for all gene family members.

## Acknowledgements

We would like to thank Honour McCann, Oliver Weichenrieder, and two anonymous reviewers for helpful comments.

## Authors’ contributions

Conceptualization, C.R.; Investigation, Z.L., C.R.; Data Curation, Z.L., C.R.; Formal Analysis, Z.L., C.R.; Visualization, C.R.; Writing –Original Draft, C.R.; Writing –Review & Editing, Z.L.,C.R.; Project Administration, C.R., Supervision, C.R.

## Data Availability Statement

All data sets used in this manuscript are publicly available on WormBase ParaSite or NCBI Genbank.

