## Supplemental Material - Figures S1-S3 for "Did *Caenorhabditis* nematodes recycle transposons to fight pathogens?"

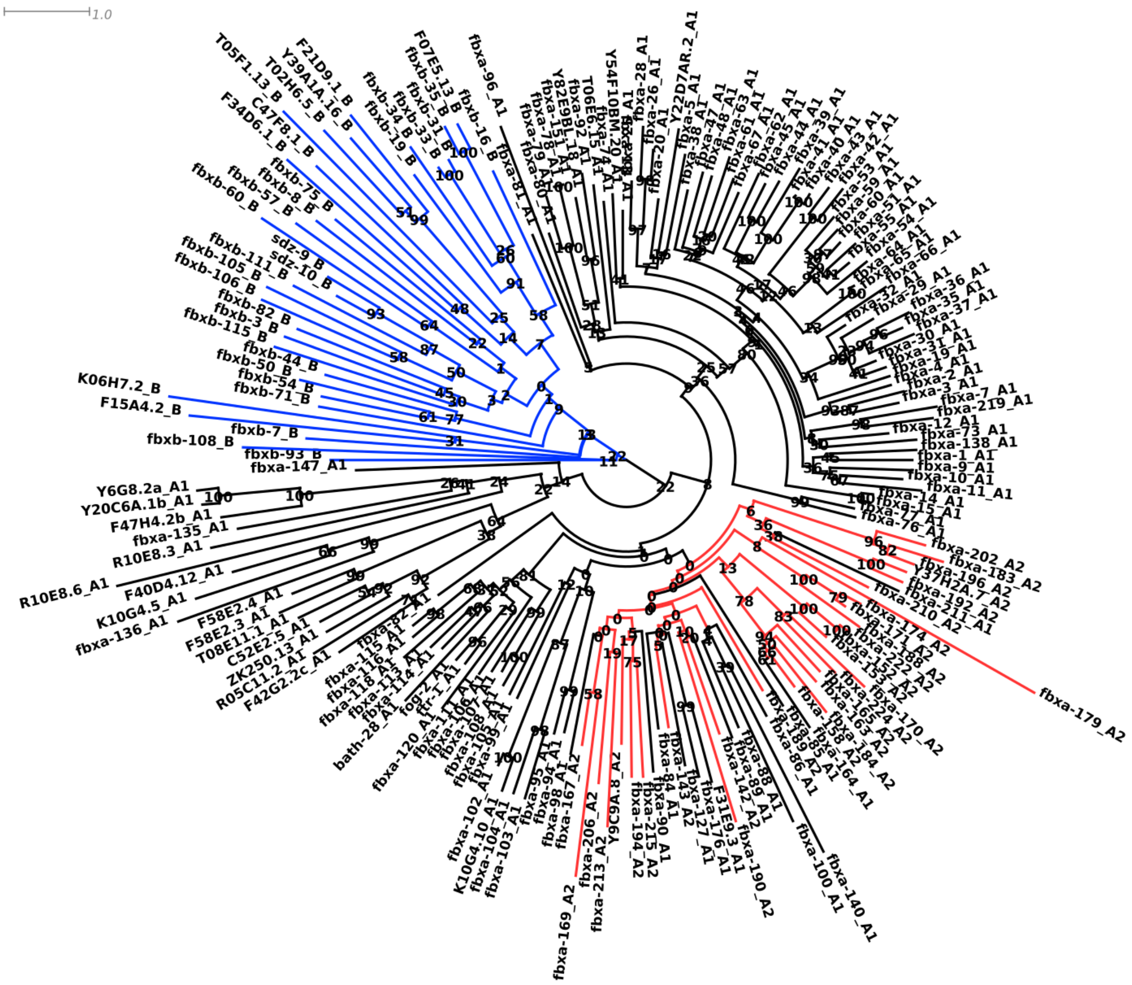


**Supplementary Figure S1.** Phylogenetic tree of *C. elegans* F-box genes with branch lengths.

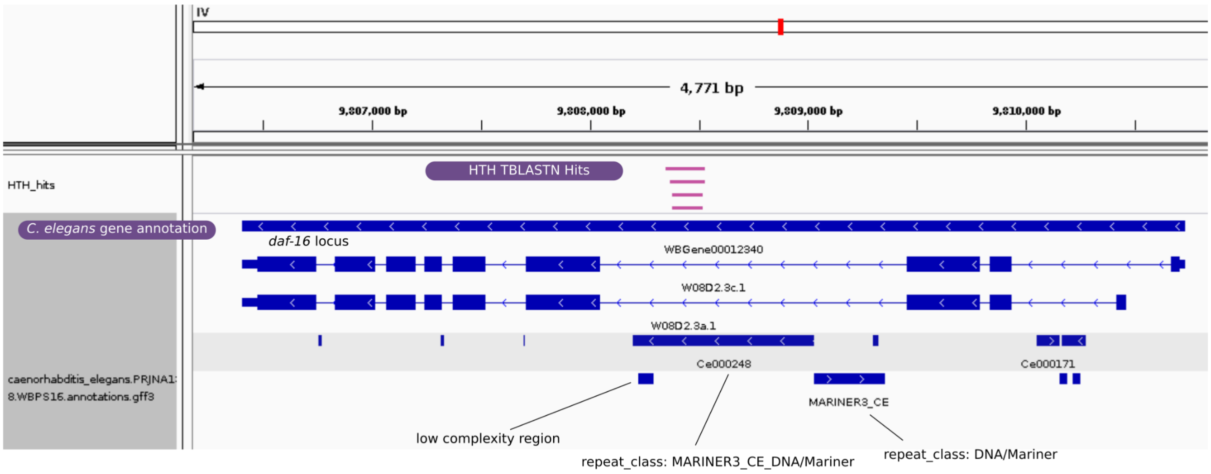


**Supplementary Figure S2.** Genomic locus of the Mariner element in *C. elegans* that represents closest relative to the HTH domains in F-box genes of the A2 family. Gene annotations and genomic coordinates of TBLASTN hits were visualized in the Integrative Genomics Viewer. The screenshot shows a 4.8kb region spanning the *daf-16* locus. The *C. elegans* gene *daf-16* has two isoforms. The largest intron contains sequences that derive from a Mariner transposon.

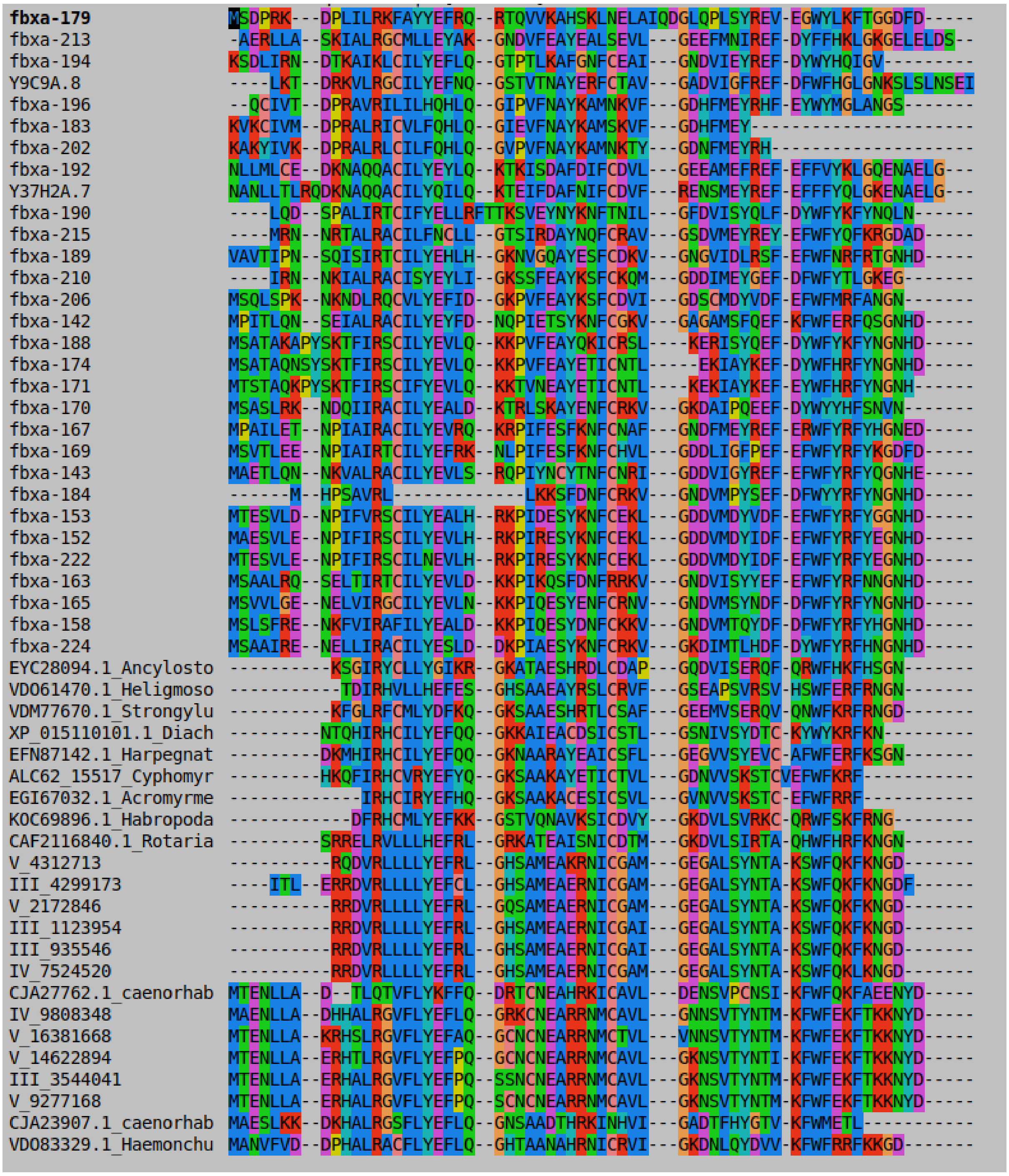


**Supplementary Figure S3. Alignment of HTH domains.** The multiple sequence alignment of HTH domains from diverse taxa was generated by the MUSCLE tool and visualized with seaview (default options).
